# Physiologically realistic gamma activity produced *in silico* by weakening the PING attractor state

**DOI:** 10.64898/2026.02.19.706788

**Authors:** Scott Rich, Tarun Shriram, Steven A. Prescott

**Affiliations:** Department of Physiology and Neurobiology, University of Connecticut, Storrs, CT, USA; Departments of Mathematics and Biomedical Engineering and Institute for Brain and Cognitive Sciences, University of Connecticut, Storrs, CT, USA; Neurosciences and Mental Health, The Hospital for Sick Children, Toronto, ON, Canada; Institute of Biomedical Engineering, University of Toronto, Toronto, ON, Canada; Department of Physiology, University of Toronto, Toronto, ON, Canada; Dept of Physiology and Pharmacology, University of Calgary, AB, Canada; Hotchkiss Brain Institute, University of Calgary, AB, Canada

## Abstract

Network oscillations in the gamma (30-80 Hz) frequency range are implicated in many vital neurological functions. The seminal Pyramidal Interneuron Network Gamma (PING) mechanism produces gamma oscillations *in silico* with dynamics associated with a strongly attracting limit cycle. However, this activity is stronger and more stable than what is observed during *in vivo* gamma oscillations, which are transient and driven by sparse spiking activity. Here we describe multiple biophysically motivated changes degenerately driving PING-motivated networks to produce more realistic gamma activity. Increased heterogeneity, synapse-like noise, and a depolarized chloride reversal potential each destabilize the attractor represented by PING-driven activity: minor alterations disrupt the unrealistic organization of idealized PING oscillations, while larger changes entirely prevent this overly-synchronous network activity. This allows new dynamics to emerge: spontaneous and transient increases in gamma-band oscillatory power with excitatory cells only weakly entrained to the population rhythm. These features, better approximating the *in vivo* reality, mirror a system’s oscillatory return to a stable focus following noise-induced perturbations. In fact, dynamical systems with both a stable focus and limit cycle are associated with Hopf bifurcations, which have been characterized in models of reciprocally connected excitatory and inhibitory neuronal populations as required for PING. We therefore propose a revised explanation for physiologically-realistic gamma activity that retains the key elements of the PING mechanism—strong reciprocal connection between excitatory and inhibitory neurons—but where biophysical phenomena bias the system towards damped oscillations around a weakly stable focus rather than a stable limit cycle.

**Significance Statement:** The mechanisms underlying network oscillations at gamma frequencies have long been a focus of computational neuroscience. This research has produced idealized mechanisms yielding highly synchronous, active, and stable oscillations; however, *in vivo* gamma rhythms are transient with sparse neuronal spiking. Here, we illustrate minimal, biophysically relevant adjustments to established Pyramidal Interneuron Network Gamma (PING) networks that bridge a major divide between these traditional *in silico* systems and the experimental reality. Uncorrelated noisy input and a depolarized chloride reversal potential disrupt stereotyped PING rhythms and promote more realistic transient increased gamma power in network activity. This important step forward in the modeling of gamma oscillations illustrates how including biophysical detail can promote more realistic activity in computational systems.

## Introduction

The brain’s capacity to produce rhythmic electrical activity is one of its most salient and well-studied features [1, 2]. Rhythms in the gamma frequency range (typically 30-80 Hz) are particularly ubiquitous [3–6], with research suggesting they serve a role in vital brain functions including learning [7], memory [8], and attention [9]. Given the perceived importance of this oscillatory activity, computational and theoretical neuroscientists have worked for decades to articulate viable mechanisms underlying oscillatory activity in networks of neurons [10, 11]. The Pyramidal Interneuron Network Gamma (PING) mechanism, which explains how strong reciprocal connectivity between excitatory (E) and inhibitory (I) neuronal populations can drive stable network oscillations at gamma frequencies [12–15], represents a significant success in this pursuit. One notable strength of this mechanism is its basis in foundational mathematical theory, specifically the presence of stable limit cycles in simplified systems with PING-motivated network topologies [16]. Dynamical systems techniques have been used to further delineate the genesis of these rhythms and their stability over time [17, 18].

Various computational studies have dissected whether the PING mechanism still applies as the idealized requirements of its original presentation are relaxed. This research has centered on the simplified question of how much biophysical realism can be tolerated before conspicuous PING oscillations “break” and asynchronous activity becomes the system’s default state. Such work has analyzed the effects of sparse and heterogeneous connectivity [19, 20], neuron models with different excitability types [21, 22] and forms of adaptation [23–25], the implementation of multiple inhibitory populations [22], and the addition of noise [17].

Indeed, as is the norm in computational modeling, the networks used in the earliest PING studies were necessarily an abstraction of the complicated biological reality [26]. Traditional PING rhythms are correspondingly idealized, consisting of highly active and synchronous bursts of excitatory cells that will continue indefinitely without an external perturbation to the system [19,20,22,23]. In contrast, *in vivo* gamma oscillations are transient [3], existing on the order of 50-100 ms [27]. They are also sparsely active [3, 28], typically identified by moderate increases in local field potential (LFP) gamma power [6, 29] rather than synchronized neuronal spiking of an entire microcircuit [4]. In a step towards capturing these features, E-I networks with heterogeneous neuronal excitability and connectivity produce gamma oscillations without every neuron participating in each gamma cycle; however, such systems are still unrealistically stable over time [22, 23]. Despite further advances from contemporary studies of gamma rhythmicity [30], a mechanistic explanation for the transience of *in vivo* gamma rhythms remains elusive.

Here, we ask whether *in silico* systems better capturing these important nuances of *in vivo* gamma oscillations might arise from enhancing the biophysical realism of PING-motivated networks. We focus primarily on two phenomena motivated by the *in vivo* reality: uncorrelated external noisy input (approximating independent synaptic-like input to each neuron) and shifts in the chloride reversal potential (which dictates the reversal potential of GABAergic signaling, *E*_GABA_ [31]). Noisy brain activity is well-characterized experimentally and is often implemented in computational models as a phenomenological implementation of inputs from other brain regions, with many studies proposing a functional role for noise in cognitive functions [32–35]. In contrast, there is a dearth of computational study of *E*_GABA_’s role in physiological oscillatory activity [36] despite evidence that altered *E*_GABA_ contributes to neurological disorders typified by abnormal oscillatory activity, including epilepsy [37–39], chronic pain [40, 41], and autism spectrum disorder [42]. However, *E*_GABA_ does not only change under pathological conditions, as it notoriously shifts during development [43] and in the mature brain varies across brain regions [37], individual neurons [39], and dynamical states [44–47].

We begin with a PING-motivated network with heterogeneous neuronal excitability and connectivity, a step away from the hyper-idealized setting of the original PING mechanism. This example network exhibits PING-driven oscillations representing a “strong attractor”: large-amplitude perturbations are required to disrupt these highly stereotyped network rhythms and yield asynchrony. Amplifying the effects of heterogeneity, adding noise, and depolarizing *E*_GABA_ all weaken this attractor. When PING-like rhythms persist, noise additionally disrupts the unrealistically consistent cell-to-(gamma) cycle entrainment of these classical oscillations. When PING-like rhythms are disrupted entirely, transient (on the order of 100s of ms) increases in the gamma band oscillatory power of network activity arise instead, more realistically approximating *in vivo* gamma activity.

These gamma events more closely resemble rhythms driven by a damped oscillation around a stable focus rather than the stable limit cycle traditionally associated with PING [16]. Indeed, similar dynamics arise from models of reciprocally connected E and I populations [48] with a stable focus. This is also true in abstract systems, often associated with a Hopf bifurcation [49], containing both a stable limit cycle and focus. Through analogy to these dynamical systems, we articulate a new theory of gamma rhythmicity in which sources of biophysical realism (including the heterogeneity, noisy external input, and depolarized *E*_GABA_ implemented here) do not “break” a system’s capacity for gamma oscillations as previously implied [17]. Instead, they merely weaken the limit cycle attractor represented by traditional PING, allowing damped oscillations (triggered by both external and internal noisiness) around the co-existing stable focus to arise. This new perspective reconciles many inconsistencies between the theoretical underpinnings and biophysical reality of gamma rhythmicity, including the brain’s capacity for both physiological, sparsely active and pathological, hyper-active and -synchronous [50] oscillations.

## Results

### Traditional PING-driven rhythms are strongly attracting with deterministic spiking

We begin with a system shown in previous work [22] to exhibit “weak PING” [51] rhythmicity: persistent network-level oscillations in which not every E neuron participates in every cycle of network activity. This less regimented excitatory activity—relative to the original PING rhythms—is driven primarily by sparse connectivity (50% inter-connectivity between E and I populations and 30% intra-connectivity within the I population) and heterogeneous neuronal excitability (implemented through variability in the external input *I*_*ext*_ to each neuron; see Materials and Methods). As illustrated in Figure 1**A**, cycles reliably alternate between containing a larger or smaller fraction of active E neurons. Indeed, despite concessions to the biological reality, these rhythms remain overly idealized [3, 28]: spiking is consistent within these alternating cycles, as illustrated by an example E neuron that exhibits perfect 1:2 cell-to-cycle entrainment highlighted in the bottom panel. Furthermore, these rhythms will continue indefinitely, another discrepancy from the biological reality [3, 27] predicted by previous analytical [17, 18] and computational [19, 20, 22, 23] studies of PING. We now ask whether this exemplar PING-driven rhythm also exhibits characteristics of a strong attractor—a stable solution only disrupted by large perturbations [52].

**Figure 1.**
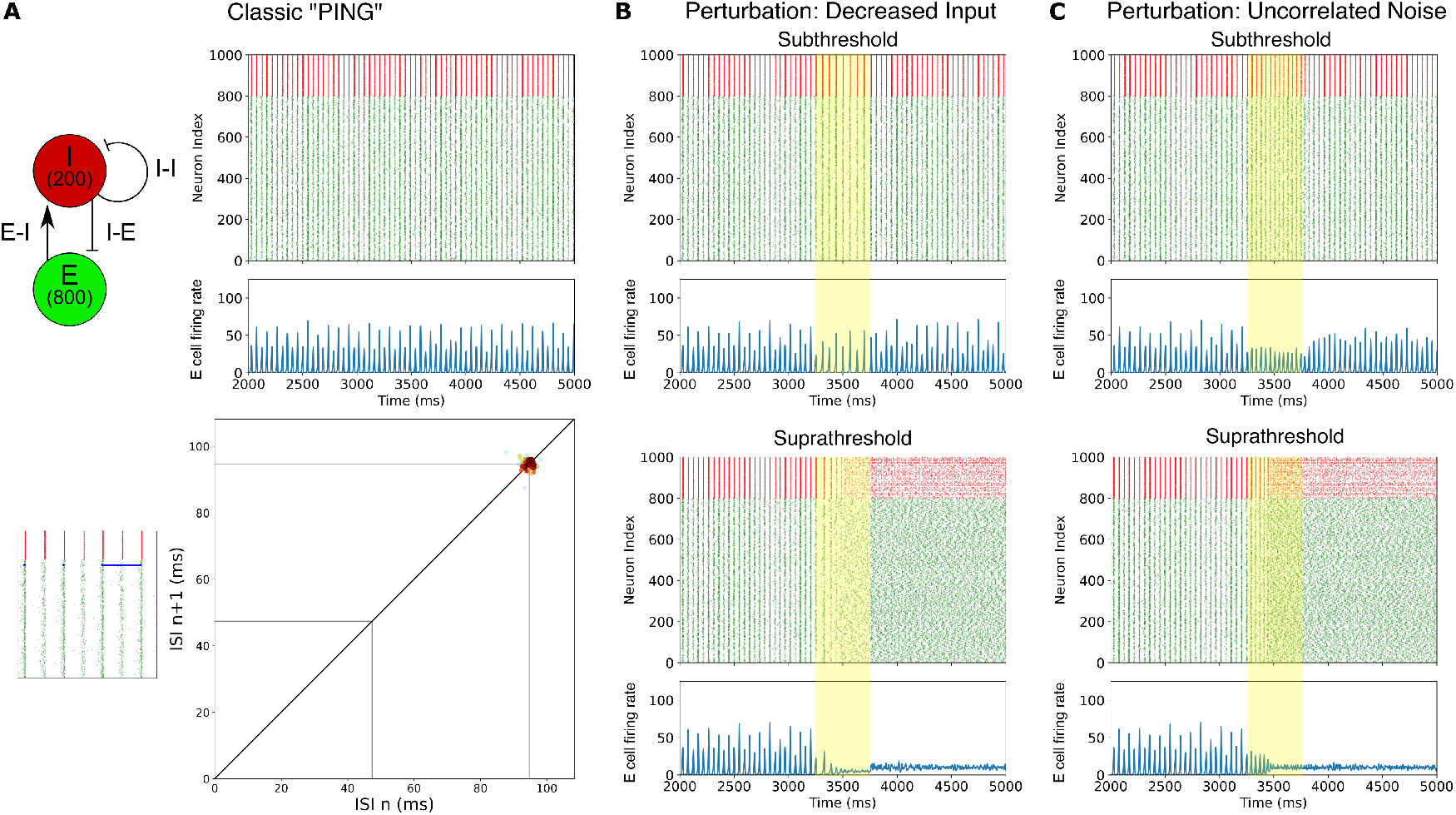
Stable “weak PING” rhythmicity represents a “strong attractor” state. **A**: Weak PING rhythms exhibited by the E-I network with chosen network topology (see Materials and Methods, circuit diagramed in top-left) are stable over time—visualized via a raster plot with excitatory cells in green and inhibitory cells in red and a continuous firing rate histogram of excitatory activity (top-right). Cell-to-cycle entrainment is consistent in these rhythms—visualized via a Pointcaré return map of an example neuron’s interspike intervals (bottom-right). Stronger “hot” colors represent the most recent events, while fainter “cold” colors represent earlier events over a 20,000 ms simulation. The raster plot snapshot on left highlights the chosen cell in blue. In this example, the chosen cell’s inter-spike intervals (ISIs) are consistently double the network oscillation’s period (integer multiples of this period are indicated via gray lines), indicative of consistent 1:2 cell-to-cycle entrainment. **B**: Perturbations lowering each neuron’s external input for 500 ms (illustrated by yellow shading) must be extreme to disrupt rhythmicity (top: halved external input; bottom: quartered external input). **C**: Perturbations adding uncorrelated Ornstein-Uhlenbeck noise for 500 ms must be extreme to disrupt rhythmicity (top: *SD*_*V*_ increased from 0 mV to 1.842 mV; bottom: *SD*_*V*_ increased from 0 mV to 2.234 mV). Raster plots and continuous firing rate histograms are as in Panel **A**.

Two types of perturbations were used to qualitatively assess the strength of this oscillation’s attractor state. First, the mean external input to each neuron was decreased for 500 ms long after the rhythm was established. The oscillation was not affected by reducing the external input by half (Figure 1**B**, top), with a larger reduction of 75 percent or more required to consistently disrupt rhythmicity (Figure 1**B**, bottom). The second perturbation was the addition of 500 ms of uncorrelated noisy input. The system was resilient to noise that would cause an isolated neuron’s subthreshold voltage to vary with a standard deviation of 1.842 mV (hereafter referred to as the *SD*_*V*_ and reported in lieu of the raw noise amplitude; see Materials and Methods), with consistent oscillations consistently disrupted by noise with a *SD*_*V*_ of 2.234 mV (Figure 1**C**). While previous studies have shown that insufficient external input and excessive noise prevent the development of stable PING rhythms [17, 53], these results are primarily contextualized in terms of the oscillations’ *development* rather than as a *perturbation* to existing rhythmicity. From this new perspective, these preliminary explorations affirm that this example PING-driven rhythm satisfies key characteristics of a strong attractor: its dynamics are stable over time in the absence of perturbation, and these stable dynamics are disrupted only by large perturbations. Thus, we will hereafter refer to this oscillation as the “PING-attractor state.” An additional notable characteristic seemingly dictated by the strength of this attractor state is that individual neuronal spiking appears deterministic given the consistent cell-to-cycle entrainment (Figure 1**A**, bottom panel).

### External noise and depolarized *E*_GABA_ weaken the PING-attractor state and promote probabilistic spiking

As discussed previously, the defining characteristics of the PING-attractor state—a strong attractor with each cell exhibiting consistent cell-to-cycle entrainment—do not correspond with experimentally-characterized gamma events that are transient with sparse E cell spiking [3, 27, 28]. Motivated by the ability of high-amplitude uncorrelated noise to force the system out of the PING-attractor state (Figure 1**C**) as well as the known variability of *E*_GABA_ experimentally [44–47], we asked whether these sources of biophysical realism typically omitted in abstract PING studies can promote more physiologically-realistic activity without requiring altered network connectivity.

We first examined the impact of varying *E*_GABA_ without external noise. An example with *E*_GABA_= −69 mV, as opposed to the −75 mV value conventionally used in previous studies of PING rhythms [22, 23], is highlighted in Figure 2**A**. Only the value of *E*_GABA_ is changed relative to simulations in Figure 1**A**, and this new value remains well within the physiologically-relevant range [44–47]. Nonetheless, this small change in *E*_GABA_ yields antithetical network dynamics: over the course of a 20,000 ms simulation, the network becomes asynchronous almost immediately (first panel), and then transitions into and out of clear rhythmicity without any external perturbation (second and third panels). This subtle change in *E*_GABA_ therefore clearly disrupts the strength of the PING-attractor state. The fact that such stochastic dynamics can arise without external noisy input is due primarily to the existence of intrinsic noisiness in the system [54], realized through heterogeneous neuronal excitability and connectivity (see Materials and Methods).

**Figure 2.**
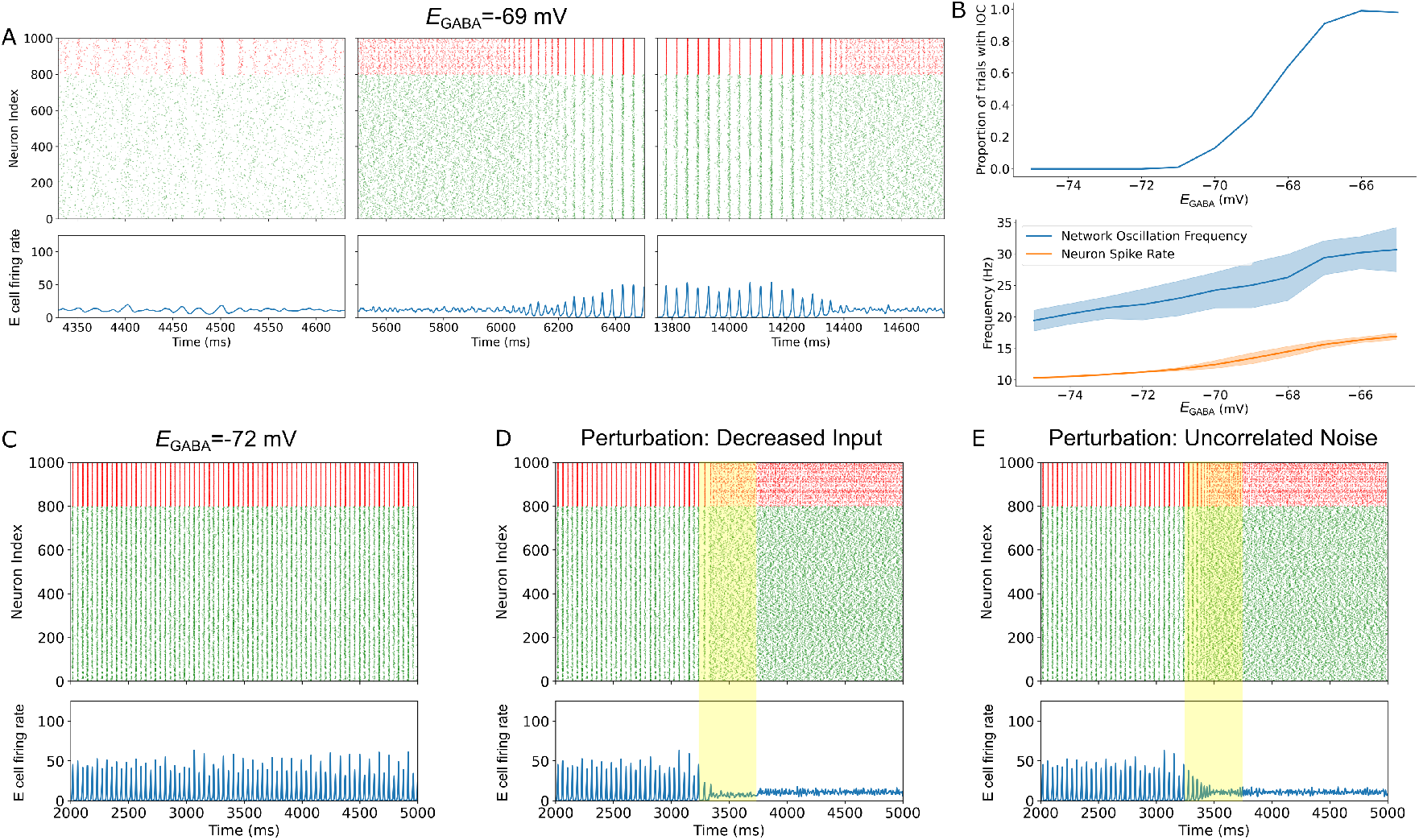
Depolarized *E*_GABA_ weakens the PING-attractor state. **A**: Example simulation with *E*_GABA_=−69 mV that spontaneously transitions into and out of PING-like oscillatory activity. **B**: Top: Proportion of 100 independent simulations that exhibit inconsistent oscillatory activity (IOC: simulations with transitions into and/or out of the oscillation, of which Panel **A** is an example) over the course of a 20000 ms simulation as a function of *E*_GABA_. Bottom: Network oscillation frequency and neuron spike rate as a function of *E*_GABA_, with shaded region representing +/-one SD. **C**: An example simulation with *E*_GABA_=−72 mV exhibiting consistent oscillatory activity (no such simulations exhibit inconsistent oscillatory activity as evidenced in Panel **B**). **D-E**: Perturbations that were insufficient to disrupt oscillatory activity with default *E*_GABA_ (see Figure 1**B-C**)— halving each neuron’s external input for 500 ms in Panel **D** and increasing the *SD*_*V*_ from 0 mV to 1.842 mV for 500 ms in Panel **E**—are enough to disrupt oscillations with this depolarized *E*_GABA_=−72 mV. Visualizations in Panels **A** and **C-E** as described in Figure 1.

We term any breakdown of the PING-attractor state—that is, any simulation in which conspicuous oscillatory activity does not persist for the entire 20,000 ms—an iteration with “inconsistent oscillatory activity.” The proportion of 100 independent simulations exhibiting inconsistent oscillatory activity is quantified in Figure 2**B**, showcasing the sensitivity of the strength of the PING-attractor state to *E*_GABA_: the PING-attractor state remains perfectly strong (i.e., no breakdown of conspicuous oscillatory activity) at *E*_GABA_=−72 mV but is extremely fragile at *E*_GABA_=−67 mV. As expected, network activity (both in the frequency of network oscillatory activity when it exists and in mean neuronal spike rates) increases as a function of depolarizing *E*_GABA_ (Figure 2**B**). Clearly, physiologically relevant fluctuations in *E*_GABA_ significantly influence the strength of the PING-attractor state, a phenomenon which is likely to influence gamma rhythmicity *in vivo*.

The impact of depolarized *E*_GABA_ also manifests in a more subtle fashion. While for *E*_GABA_=−72 mV the PING-attractor state continues to dominate network activity, the attractor becomes weaker: rhythmicity (Figure 2**C**) can be disrupted by smaller perturbations, whether in decreased input (Figure 2**D**) or uncorrelated noise (Figure 2**E**). Therefore, even small changes in *E*_GABA_ render the system more vulnerable to perturbations disrupting the overly-idealized oscillatory activity of the PING-attractor state.

The main effect of depolarized *E*_GABA_ is to weaken inhibition. However, we were unable to reproduce the dynamics caused by depolarized *E*_GABA_ merely by decreasing the inhibitory synaptic strengths (*g*_*syn*_ values for I-I and I-E synapses; see Materials and Methods), as illustrated in Supplementary Figure S1. Indeed, these two means of disrupting inhibitory synaptic signaling act in distinct fashions mathematically: decreasing *g*_*syn*_ decreases the strength of inhibitory synapses in a constant manner, while depolarized *E*_GABA_ weakens the synaptic current’s driving force dependent on the post-synaptic voltage and thus varies over time. This finding points to a potentially unique role for *E*_GABA_ in the stability of oscillatory activity in neuronal microcircuits that has not been previously reported.

We next jointly analyzed the effects of varying *E*_GABA_ and noisy input (see Materials and Methods) on the strength of the PING-attractor state. The effects of noise on the likelihood of our previously defined “inconsistent oscillatory activity” is illustrated in Figure 3**A**: low amplitude noise leads to a right-shifted but steeper sigmoid, while high amplitude noise leads to a strong leftward shift until stable oscillatory activity is never observed. Noise levels have minimal influence over the population oscillatory frequency (Figure 3**B**) and mean neuronal spike rate (Figure 3**C**). While these analyses highlight complex interactions between noise and *E*_GABA_, a key unifying feature is that both low- and high-amplitude noisy input result in the complete disruption of consistent oscillatory activity for less depolarized values of *E*_GABA_ than in the absence of noise.

**Figure 3.**
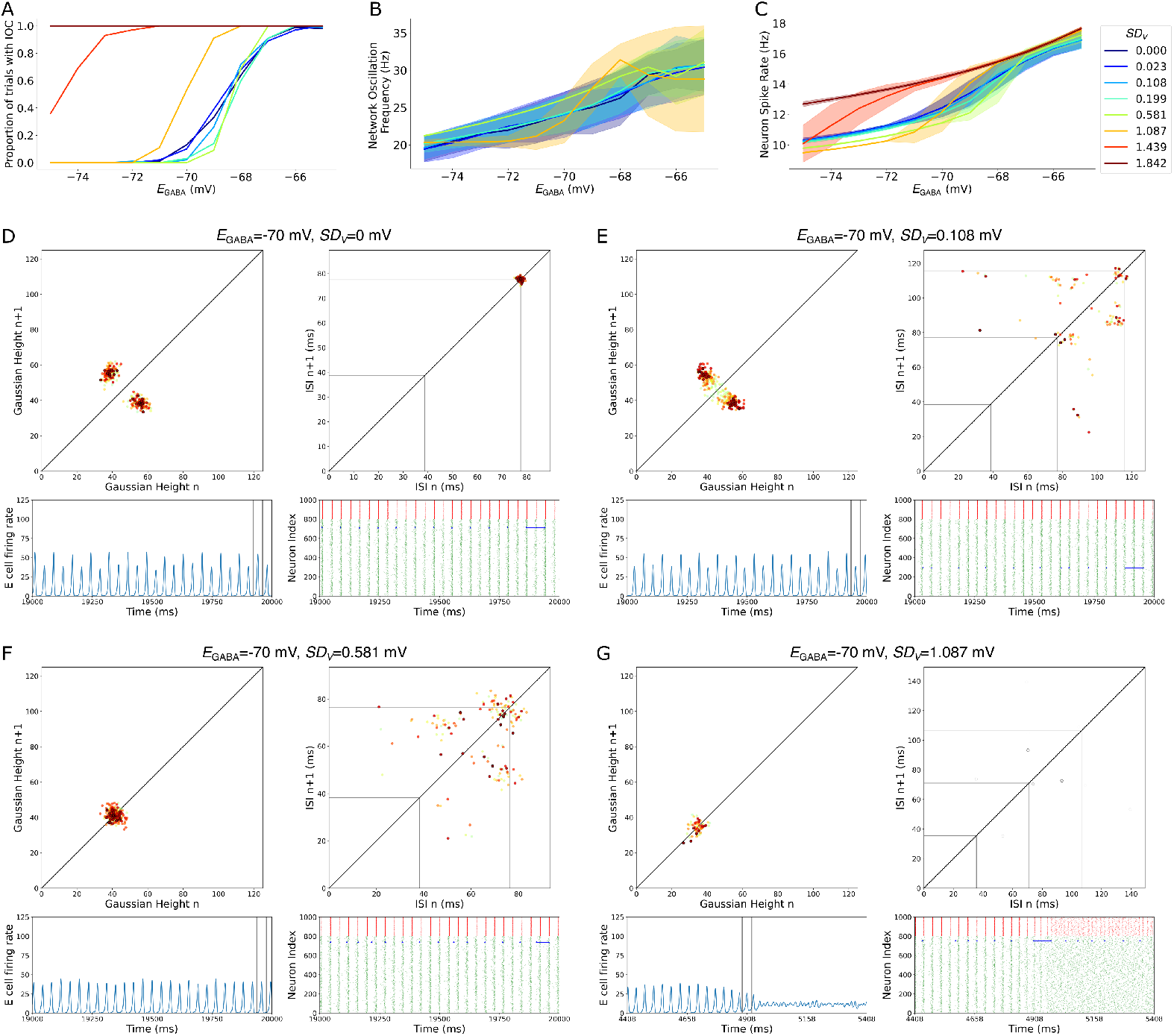
Uncorrelated noisy input disrupts cell-to-cycle entrainment at low amplitudes and disrupts PING-attractor state at high amplitudes. **A**: Low amplitude noise prevents state transitions at moderately depolarized *E*_GABA_ but causes such transitions uniformly at more depolarized *E*_GABA_ values (i.e., a rightward shifted but higher gain sigmoid). High amplitude noise leads such transitions to arise for less depolarized *E*_GABA_ (i.e., a leftward shifted sigmoid), with sufficiently high amplitude noise preventing stable oscillations entirely. **B-C**: Network oscillation frequency (Panel **B**; curves for two highest amplitude noise values omitted due to insufficient network oscillations) and neuron spike rate (Panel **C**) as a function of *E*_GABA_ and noise amplitude (plotted as mean *±* SD) are minimally affected by low-to-moderate amplitude noisy input. **D-G**: Example dynamics of systems with *E*_GABA_=−70 mV with various levels of noisy input. Without noise (Panel **D**; full dynamics in Supplementary Movie S1), network dynamics mimic traditional PING rhythms (see Figure 1) with stable alternating network activity (left panels) and consistent cell-to-cycle entrainment (right panels). Low amplitude noise (Panel **E**; full dynamics in Supplementary Movie S2) causes variable cell-to-cycle entrainment (right panels) while preserving the alternating network activity (left panels). Higher amplitude noise (Panel **F**; full dynamics in Supplementary Movie S3) promotes further variability in cell-to-cycle entrainment (right panels), which manifests in more uniform, but less active, bursts in network activity (left panels). Sufficiently high amplitude noise (Panel **G**; full dynamics in Supplementary Movie S4) creates network bursts with few active neurons (left panels), leading the network to quickly decay to asynchrony. Figures show Pointcaré return maps (top left) of continuous firing rate histogram peaks (bottom left), and Pointcaré return maps (top right) of an individual cell’s inter-spike intervals (highlighted in blue on the raster plot in the bottom right), with gray lines representing integer multiples of the network oscillatory period. In return maps, stronger “hot” colors represent the most recent events, while fainter “cold” colors represent earlier events; events towards the end of the simulation are emphasized in the static images, while full dynamics can be seen in the Supplementary Movies.

More conspicuous is the effect of noise on cell-to-cycle entrainment, illustrated for *E*_GABA_=−70 mV in Figure 3**D-G**. Without noise (Figure 3**D**), the system organizes itself into stable alternating cycles containing a low or high proportion of active E neurons with consistent cell-to-cycle entrainment, similar to the default *E*_GABA_=−75 mV in Figure 1. With low-amplitude noise the stable alternating cycles persist (Figure 3**E**, left panels) but cell-to-cycle entrainment is disrupted (right panels). Cell-to-cycle entrainment is further disrupted by higher amplitude noise (Figure 3**F**, right panels), which also disrupts the stable alternating cycles; instead, a smaller but similar number of neurons spike on each cycle (Figure 3**F**, left panels). Finally, with sufficiently strong noise early oscillations have highly irregular cell-to-cycle entrainment and uniformly small participation in each cycle, leading oscillatory activity to decay entirely (Figure 3**G**).

Thus, the effects of noise on the PING-attractor state can manifest differently than those of depolarized *E*_GABA_, particularly for low-to-moderate amplitude noise that does not entirely disrupt conspicuous, PING-like oscillations. While sufficiently depolarized *E*_GABA_ primarily causes inconsistent oscillatory activity (see the case of *E*_GABA_=−69 mV in Figure 2**A**), uncorrelated noise seems to alter the spiking pattern underlying those oscillations. This is evidenced in the examples highlighted in Figure 3: cell-to-cycle entrainment is consistent for *E*_GABA_=−70 mV with no noise (1:2 for the example cell in Figure 3**D**), while each example with noise (Figure 3**E-G**) contains neurons with variable ISIs indicative of variable cell-to-cycle entrainment. However, in these examples with *E*_GABA_=−70 mV, conspicuous population-level oscillations persist for the entire 20,000 ms simulation for all but the largest amplitude noise. Considering the similarity in neuron spike rate for all but the largest amplitude noise (Figure 3**C**), the primary effect of noise appears to be the promotion of less deterministic participation of neurons within each cycle, which typifies physiological gamma rhythms but is absent in traditional PING oscillations.

### Systems biased away from the PING-attractor state exhibit physiologically-realistic gamma events

A paramount finding from varying *E*_GABA_ was the network’s dynamics in periods without conspicuous oscillations. An example is included in the leftmost panel of Figure 2**A**: at around 4500 ms clear bursts of I cell spiking and fainter changes in E cell spiking arise but persist for no more than 100 ms. We hypothesized that such dynamics arise when the PING-attractor state is too weak to maintain oscillations with the characteristics of a stable limit cycle.

To test this hypothesis, we first focused our parameter space: we varied just the noise amplitudes while maintaining the default *E*_GABA_=−75 mV, instead further weakening the PING-attractor state by adjusting the system’s initial conditions. Indeed, highly active initial conditions are ideal for PING-driven activity to arise given the need for sufficient E cell spiking [53]. Instead, we designed “sparsely active” initial conditions (illustrated by example raster plot in Figure 4**A**; see Materials and Methods) which weaken the PING-attractor state by amplifying the effects of the system’s inherent heterogeneities (neuronal excitability and connectivity; see Materials and Methods). This allows effects of heterogeneity to fully manifest before being drowned out by network synchrony. Adding noisy input further weakened this attractor as illustrated in Figure 4**A**: the level of tonic external drive needed to consistently yield stable oscillatory activity (in these scenarios, once PING-like oscillations arise they persist for the remainder of the simulation) increases notably as a function of noise intensity.

**Figure 4.**
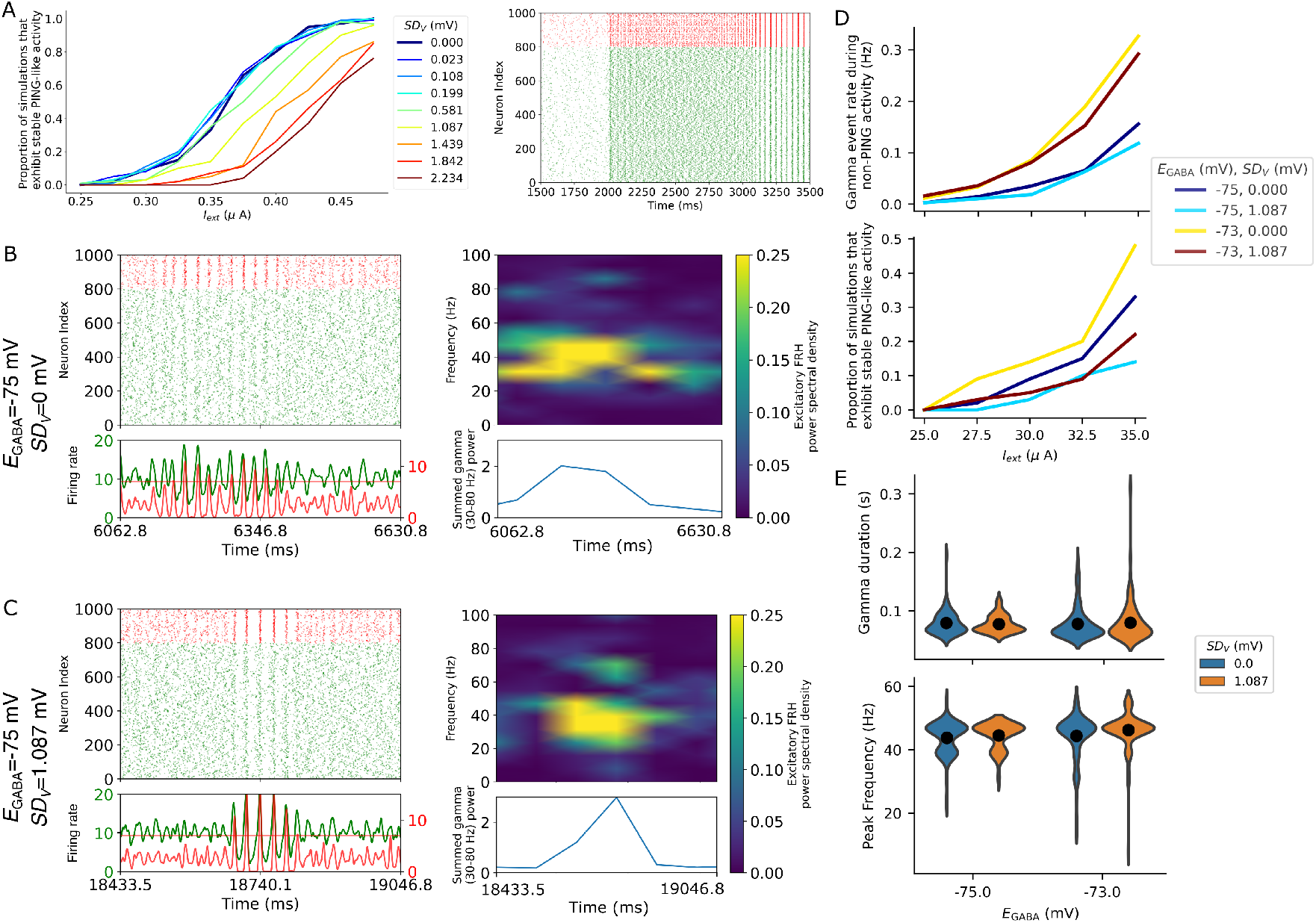
Destabilizing the PING-attractor state promotes more physiologically realistic gamma activity. **A**: Proportion of 100 independent simulations that eventually exhibit stable, PING-like oscillatory activity as a function of external input and noise level following sparsely active initial conditions, exemplified by the raster plot (*I*_*ext*_=.40 *µ*A, no noise) on right. **B-C**: Examples of transient, weak increases in gamma activity at default *E*_GABA_ and *I*_*ext*_=.30 *µ*A without (Panel **B**) and with (Panel **C**) noise, as evidenced in the raster plot (top left), continuous firing rate histogram (bottom left), and spectrogram of the excitatory firing rate histogram (right). **D**: Top: rate of detected gamma events (Hz; averaged over 100 independent simulations) during non-PING activity for four combinations of noise and *E*_GABA_. Rates are significantly different (two-sample t-test, p*<*0.05) across all *I*_*ext*_ for any comparison between different *E*_GABA_ values. Rates for *E*_GABA_=−75 mV with and without noise are significantly different only for *I*_*ext*_=0.30 *µ*A and 0.35 *µ*A. Rates for *E*_GABA_=−73 mV with and without noise are significantly different only for *I*_*ext*_=0.325 *µ*A. Bottom: proportion of 100 independent simulations that eventually exhibit stable, PING-like oscillatory activity (as in Panel **A**). Proportions are significantly different (two-sample t-test, p*<*0.05) between *E*_GABA_=−75 mV with and without noise only for *I*_*ext*_=0.35 *µ*A; between *E*_GABA_=−75 and −73 mV without noise for *I*_*ext*_=.275 *µ*A and *I*_*ext*_=0.35 *µ*A; for *E*_GABA_=−73 mV with and without noise for *I*_*ext*_ ≥0.30 *µ*A; for *E*_GABA_=−75 with noise and *E*_GABA_=−73 without noise for *I*_*ext*_ ≥27.5 mV. All analyses here are performed for a narrower range of *I*_*ext*_ than in Panel **A**, centered on the exemplar *I*_*ext*_=.30 *µ*A yielding Panels **B** and **C. E**: Features of gamma events at *I*_*ext*_=.30 *µ*A (averaged over 100 independent simulations) with and without noise at both default and slightly depolarized *E*_GABA_. Differences between the mean gamma duration (top) are not statistically significant (two-sample t-test, p*>*.64 for all comparisons); differences between the mean peak frequency during gamma events (bottom) is significant (two-sample t-test, p*<*0.05) only when comparing the scenario with the highest mean peak frequency (*E*_GABA_=−73 mV and *SD*_*V*_ =1.087) to the means of scenarios without noise (both *E*_GABA_=−75 and −73 mV, *SD*_*V*_ =0).

We further examined the case of *I*_*ext*_=0.30 *µ*A, which rarely yields stable oscillations without noise. We identified transient oscillations, termed gamma events, associated with high amplitude fluctuations in I cell firing, and therefore used the I cell firing rate histogram to detect said events (see Materials and Methods). Spectral analysis of the firing rate histogram for E cells revealed that such periods are typified by notable increases in gamma-band power, both without (Figure 4**B**) and with (Figure 4**C**) noise. These events better reflect the gamma oscillations observed experimentally than any PING-related oscillation described previously in this manuscript or in the broader literature (see Introduction).

To confirm that this phenomenon is not reliant upon our default *E*_GABA_ value, we quantified the rate of these events (only considering the time spent in non-PING activity) for a slightly depolarized *E*_GABA_ = −73 mV both without noise and with a moderate-amplitude noisy input (Figure 4**D**, top). In fact, such events are more frequent with depolarized *E*_GABA_ regardless of noise. For completeness, we quantified two key features of these events, the frequency of peak spectral power and the events’ temporal duration. Both *E*_GABA_ and noise minimally affect these features: the mean gamma duration varies between 77.3 and 79.5 ms and the mean peak frequency varies between 43.8 and 46.3 Hz (well within the traditional range defining gamma frequencies) over all four conditions (Figure 4**E**). Noise also acts to decrease the likelihood that the system will transition into stable PING-like rhythms for *E*_GABA_ = −73 mV as it did for our default *E*_GABA_ = −75 mV (Figure 4**D**, bottom).

Collectively, these results illustrate that noise has a minimal effect on the rate and features of gamma events but a significant influence over the chance that PING-like dynamics arise (see testing details in Figure 4 caption). This indicates that noise’s primary role in this setting is to bias the system away from the PING-like oscillations that inherently preclude gamma events. This is relevant to the stochasticity of the dynamics illustrated by the raster plot in Figure 4**B**, which is driven not by *external* noise but rather *internal* noisiness in this system, created by heterogeneous neuronal excitability and connectivity (see Materials and Methods) and amplified by the sparsely active initial conditions. The effects of this internal noisiness become more apparent the longer the system exists in a sparsely and asynchronously firing state rather than the hyper-active and -synchronous activity typifying PING-like rhythmicity. This explains its more pronounced effects under this experimental paradigm than with our previous initial conditions predisposed towards PING-like rhythmicity from the simulation’s onset. The multiple roles that both external and internal noise play in this system will be described further in the Discussion.

### A dynamical mechanism explaining the co-existence of transient and stable oscillations

We see from these explorations that the same network topology can yield both stable and transient oscillatory activity, a phenomenon explainable via mathematical analogy. Dynamical systems existing near a subcritical Hopf bifurcation [49] commonly exhibit a stable fixed point (focus) surrounded by a stable limit cycle, with an unstable limit cycle between them defining the basins of attraction. A simple dynamical system exhibiting this stability structure, derived in polar coordinates (see Materials and Methods), is illustrated via its phase diagram in Figure 5**A**. To facilitate later comparison, we refer to the Cartesian x-coordinate as *r*_*e*_ and the Cartesian y-coordinate as *r*_*i*_, analogues for the unitless population activity variables (for excitatory and inhibitory populations, respectively) used in the seminal Wilson-Cowan equations [48]. In the bottom of Figure 5**A** we show, through numerical simulation, that a sufficient perturbation away from the stable fixed point will trigger transient oscillatory dynamics in this system; however, too large of a perturbation will force the system beyond the unstable limit cycle and cause the stable limit cycle to dominate.

**Figure 5.**
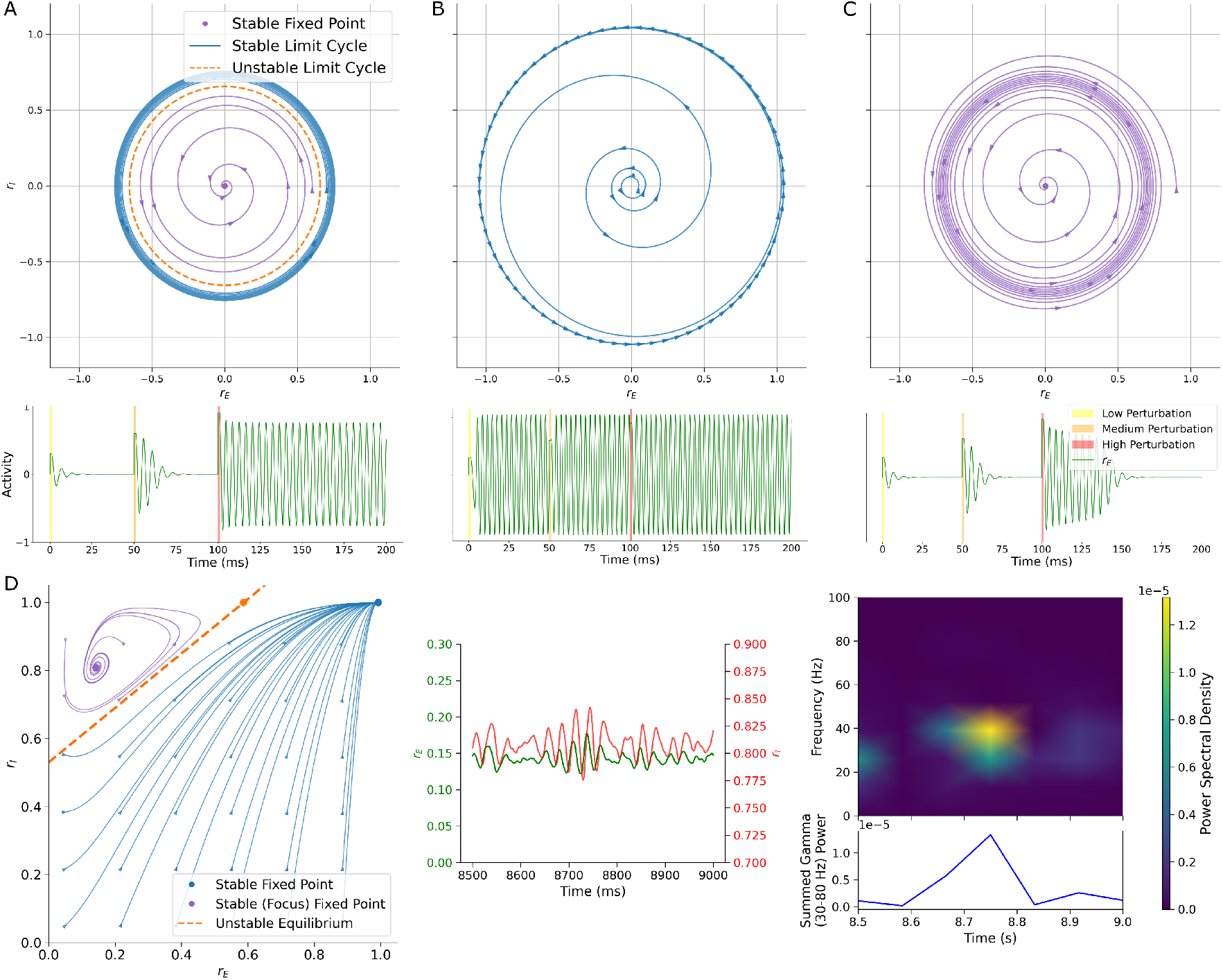
Transient oscillations emerge as dynamical systems return to a weakly stable fixed point (focus). **A:** Top: Phase portrait showing trajectories in a system arising from a subcritical Hopf bifurcation in polar coordinates (see equations in Materials and Methods). Trajectories starting inside the unstable limit cycle (dashed orange) spiral toward a stable fixed point (purple), while those starting outside spiral out to a stable limit cycle (blue). Bottom: Time series showing how perturbations of increasing amplitude (see Equation 4 in Materials and Methods) induce transient oscillatory dynamics when activity is contained by the unstable limit cycle (first two perturbations), compared to stable oscillations when the (final) perturbation pushes the system beyond the unstable limit cycle. **B:** Panels as in Panel **A**, where the system now only has a stable limit cycle which dictates dynamics regardless of perturbations. **C:** Panels as in Panel **A**, where the system now only has a stable fixed point which can yield transient (i.e., damped) oscillatory behavior with a sufficiently strong perturbation. **D:** Left: Phase portrait as in Panel **A** of a version of the Wilson-Cowan equations (see Equation 5 in Materials and Methods) characterized by two stable fixed points, one oscillatory (purple) and one not (blue), separated by an unstable trajectory (orange). Right: Subjected to Ornstein-Uhlenbeck noise, these Wilson-Cowan equations exhibit transient gamma-band oscillations with conspicuous similarities to those seen in our spiking neuronal network (Figure 4**B-C**). These arise when noise pushes the system away from the stable oscillatory fixed point before returning via a damped oscillation.

One can consider this stable limit cycle as representing the PING-attractor state. The system is always capable of exhibiting a PING-driven rhythm if it is forced into the stable limit cycle’s basin of attraction; however, the likelihood of this occurring depends critically on the location of the unstable limit cycle. The mathematical system in Figure 5**A** therefore approximates our network subject to a combination of sparsely firing initial conditions, moderate amplitude noisy input, and/or moderately depolarized *E*_GABA_, as the proximity of the unstable and stable limit cycles means the system requires a large perturbation or precisely chosen initial conditions to enter the basin of attraction of the PING-attractor state. However, the stable focus can still create transient oscillatory dynamics via damped oscillations (discussed in detail below).

In contrast, a noiseless system with default *E*_GABA_ would be akin to a system where the unstable limit cycle and the stable focus collide and cease to exist (Figure 5**B**)—the stable limit cycle is the lone attractor, making PING-like oscillations the only possible dynamical outcome. At the other extreme, a system incapable of achieving a PING-like oscillation in any scenario (e.g., systems with the lowest external drives in Figure 4**A**) would only contain the stable focus and may exhibit transient oscillatory activity depending upon the strength of said fixed point and any perturbations to the system (Figure 5**C**).

To further this analogy we directly analyzed the Wilson-Cowan equations [48], a seminal system known to exhibit PING-like oscillatory dynamics driven by the reciprocal connectivity of excitatory and inhibitory populations. In this formalism we derived a parameter set with similar stability features as the idealized system described in Figure 5**A**, albeit minus the presence of a stable limit cycle; instead, a stable focus and stable node are separated by a boundary passing through an unstable fixed point (Figure 5**D**, left). Systems with a stable focus are known to exhibit damped oscillations as the system relaxes after being perturbed away from the fixed point [55]; examples of this phenomenon are seen in Figure 5**A** and **C**. Numerically simulating this system with noisy external input—of small magnitude so that the system will not cross the unstable boundary—yields dynamics paralleling those in Figure 4: transient increases in gamma power arise without the system exhibiting idealized (high amplitude and stable) oscillations (Figure 5**D**, right).

Collectively, these mathematical explorations yield a theoretical explanation for the phenomenon illustrated in Figure 4. Approximations of the gamma events observed in our spiking microcircuit arise from damped oscillations around a stable focus in the Wilson-Cowan equations. Furthermore, dynamical systems existing near a subcritical Hopf bifurcation exhibit both a stable focus and stable limit cycle (previously associated with PING-driven rhythms [16]). Systems in which PING-like oscillations arise consistently and persist over time therefore parallel dynamical systems dominated by the basin of attraction of the stable limit cycle. Biophysical properties added to idealized PING networks—including heterogeneity, synapse-like noise, and depolarized *E*_GABA_—do not destroy the stable limit cycle, but shrink its basin of attraction so that transient gamma events arising from damped oscillations can arise in the default dynamical state.

## Discussion

Physiological gamma oscillations *in vivo* are transient [3,27] with sparsely active [3,28] and weakly synchronous [4] neuronal spiking. While the seminal PING mechanism has proven invaluable by describing how idealized gamma rhythms might arise, the dynamics it produces do not accurately reflect these experimental observations. Here, we show that dynamics arising from PING-motivated network topologies will better approximate these key nuances of gamma activity under more biophysically realistic conditions, as realized through the interacting effects of neuronal heterogeneity (amplified via sparsely active initial conditions), synapse-like independent noisy input, and depolarizing shifts in *E*_GABA_. We explain this phenomenon by analogy to dynamical systems associated with a Hopf bifurcation [49] that have both a stable focus and limit cycle.

Starting with a network exhibiting classic “weak PING” oscillations as its strong attractor state (Figure 1), we illustrate that physiologically relevant depolarizing shifts in *E*_GABA_ weaken this attractor state, with the strongest shifts causing inconsistent oscillatory activity (Figure 2). Meanwhile, uncorrelated noisy input disrupts the consistent cell-to-cycle entrainment that in part contributes to unrealistically hyperactive spiking [50] in idealized PING oscillations, with high amplitude noise disrupting conspicuous oscillations entirely (Figure 3). Sparsely active initial conditions, which ensure the full effects of the network’s heterogeneous neuronal excitability and connectivity manifest at the onset of a simulation, also make such oscillations less likely. Indeed, each of these three manipulations can be viewed as degenerately biasing the system away from overly idealized PING rhythmicity. With this attractor sufficiently weakened, transient increases in gamma-band oscillatory power arise spontaneously (Figure 4), a more realistic approximation of gamma activity observed *in vivo*.

Such dynamics are much more similar to damped oscillations around a stable focus (Figure 5) than the stable limit cycle traditionally associated with PING [16]. The fact that our E-I network topology can support both PING-like rhythms and transient gamma activity therefore indicates that its dynamics are analogous to those of a system with both a stable focus and limit cycle, a conclusion strengthened by the strong parallels between the activity of our complex system and simpler ones with this structure (Figure 5). Through the lens of dynamical systems, there are two requirements necessary for damped oscillations to be the primary driver of rhythmicity: 1) The basin of attraction of the stable limit cycle must be sufficiently small, and 2) There must be a source of noise perturbing the system away from the stable focus. Depolarized *E*_GABA_ and weakening the system’s tonic external input (under sparsely active initial conditions) contribute to satisfying the first requirement, leading to inconsistent oscillatory activity and, in extreme cases, the abolition of any PING-like rhythms. While external noisy input and the system’s inherently heterogeneous neuronal excitability and connectivity also contribute degenerately to weakening the PING-attractor state, they additionally satisfy the second requirement: this manifests both in the existence of transient gamma events and also less deterministic spiking during traditional PING rhythmicity.

While any number of additions to our PING-motivated network topology might fulfill one of these two dynamical requirements, we consciously chose processes reflecting relevant biological phenomena: noise approximating external synaptic input [56–59], depolarized *E*_GABA_ reflecting physiological settings [44–47], and initial conditions better capturing the default sparsely and asynchronously firing state of the cortex [60]. This approach follows the established tradition in computational neuroscience [17–25, 61] of building upon foundational mechanisms and idealized systems with biophysical realism to better approximate and understand the complex brain. In so doing, we are able to draw new insights regarding the functional purpose of these phenomena.

While a number of potential functions have been proposed for the inherent noisiness of the brain, it has remained challenging to reconcile this feature with the brain’s capacity for oscillatory dynamics. Indeed, previous literature has shown that noise disrupts PING-driven oscillatory activity [17], although this is not uniformly the case for all oscillatory mechanisms [62]. However, given the highly idealized nature of the original PING mechanism, it is possible that this conflict is an artificial consequence of this mechanism operating far from the biophysical reality. Our results instead indicate that noise plays a necessary role in driving physiologically-realistic gamma activity. Rather than merely “breaking” stable oscillatory dynamics [17]—i.e., destroying the dynamical system’s stable limit cycle—noise biases the system away from the limit cycle’s basin of attraction while driving transient gamma events via damped oscillations [55]. The inherent heterogeneity of our model system also manifests in “intrinsic” noise in the system that allows for such activity in the absence of external noisy input. These network features have been shown in recent theoretical work to drive variable spiking in recurrent networks [54].

Some effects of depolarized *E*_GABA_ are degenerate with those of noise, particularly under sparsely active initial conditions (Figure 4). However, *E*_GABA_ also more directly influences the strength of the PING attractor state (Figure 2): slightly depolarized *E*_GABA_ transitions a system acting as a “strong attractor” [52] into one that can allow network dynamics to stochastically shift into and out of idealized PING-like oscillations. We argue that this serves a vital physiological purpose, creating conditions more permissive of the damped oscillations that mirror physiological gamma activity in lieu of unrealistically stable rhythms.

It bears emphasizing that we here borrow terminology from dynamical systems theory primarily in an illustrative sense. While we present example systems to make our argument through analogy, these are not direct analyses of the spiking neuronal network simulated throughout this manuscript—the complex mathematical analysis required to do so is outside the scope of this paper. Instead, our arguments are based off of foundational features of dynamical systems that correspond with our observed activity patterns.

There are key distinctions between the questions asked in this study and those in previous PING-focused literature. The most rigorous study of the effects of noise on PING activity [17] focuses on how noise disrupts existing oscillations, primarily via “phase walkthrough.” While some examples of noise completely disrupting the stereotypical “bursts” of network activity are illustrated, phase walkthrough primarily results in unstable network bursting that gradually decays. The focus of this previous work is on the conditions under which idealized PING-driven rhythms might “withstand” the addition of noise. While our results confirm the general principles put forth in this previous research, our primary interest is how noise (and *E*_GABA_, a property notably not the focus of any rigorous study relative to PING rhythmicity) affects both the onset and maintenance of idealized PING oscillations and, in turn, the ability of a network to exhibit more realistic transient gamma events. Through this perspective, the tendency of noise to oppose stable PING-driven activity is recontextualized as serving a necessary role in promoting more biophysically-realistic oscillatory activity.

We also note that we consciously chose not to vary parameters determining network connectivity (i.e., the synaptic conductances and connection probabilities) to maintain a reasonably constrained parameter space; the effects of these parameters on PING rhythms have been studied in previous literature [22, 23]. We specifically chose a network exhibiting PING-like oscillations slightly below the traditional gamma frequency range so that the system was not overly idealized to the point that PING was the only possible dynamical outcome; indeed, slower PING-driven activity has been shown to be more fragile in previous work [17].

It is common in computational modeling to consider an idealized, homogeneous system in order to facilitate mechanistic understanding, a philosophy implemented by necessity in the original articulation and study of the PING mechanism [12–15]. However, the brain is inherently more complex, as notable heterogeneity is not only observed experimentally [63–69] but may be functionally relevant [69, 70]. Our results follow this trend: the inclusion of various forms of biological realism make an otherwise idealized system more complex and heterogeneous, leading to dynamics better capturing the physiological reality. As our understanding of the brain’s complexity and heterogeneity at multiple spatial scales continues to evolve, it is vital that the effects of this variability on existing computational models is understood. Indeed, as is the case here, these effects may improve the capacity for these models to capture the physiological reality.

## Materials and Methods

### Neuron Model

Model neurons implemented in this study follows the Hodgkin-Huxley, conductance-based formalism with parameters adjusted to approximate features of cortical pyramidal neurons [71, 72] and fast-spiking parvalbumin-positive (PV+) interneurons [22, 23]. The equations governing this model are:

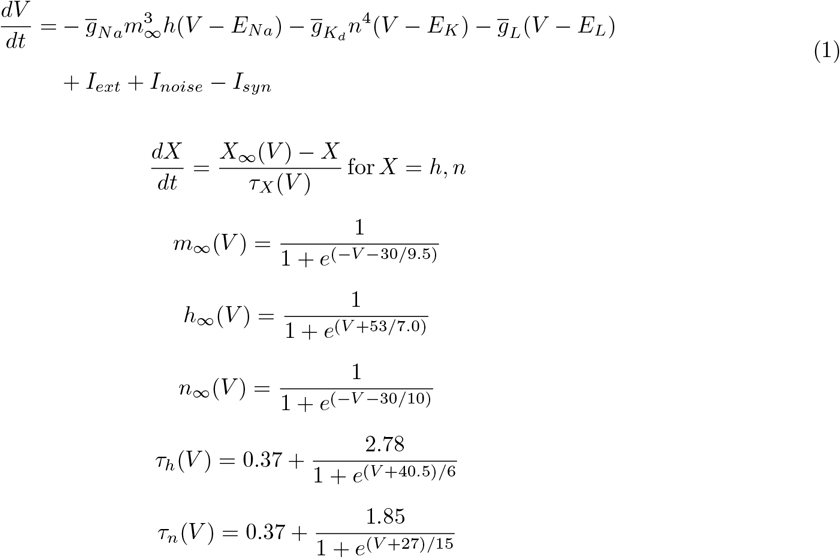

*V* represents the membrane voltage in mV, while *m, n* and *h* represent the unitless gating variables of ionic conductances. *E*_*Na*_, *E*_*K*_ and *E*_*L*_ are the reversal potentials and 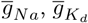 and 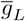 are the maximal conductance densities. *Na* represents the sodium current, *L* the leak current, and *K*_*d*_ the delayed rectifier potassium current. We omit the slow M-type potassium current in this presentation as we exclusively studied neurons in this formalism without this current active, yielding Type I excitability profiles [72]. Values for the remaining parameters are *E*_*Na*_ = 55 mV, *E*_*K*_ = −90 mV, *E*_*L*_ = −60 mV, 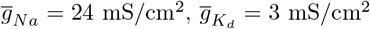, and 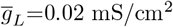. *I*_*xt*_, *I*_*noise*_, and *I*_*syn*_ are defined below.

### Network Structure

Networks studied here consist of 800 excitatory (E) and 200 inhibitory (I) neurons, matching the ratio typically used in the study of E-I networks [22, 23, 69]. *I*_*app*_ is defined differently for E and I cells to mirror traditional studies of the PING mechanism. A moderate level of cellular heterogeneity [69] is implemented by sampling *I*_*app*_ uniformly from the distribution [0.9*I*_*ext*_, 1.1*I*_*ext*_] *µ*A/cm^2^, where *I*_*ext*_ is the mean external input reported in the results. I cells receive a slight hyperpolarizing input to ensure they do not spike without excitatory input along with mild heterogeneity: for each I cell, *I*_*app*_ is sampled uniformly from the distribution [−0.21, 0.19] *µ*A/cm^2^.

Network connectivity varies in each independent simulation of a particular system. Each neuron synapses onto each other neuron probabilistically, with *p* = 0.5 for E-I and I-E connections and *p* = 0.3 for I-I connections. No E-E synapses are modeled. The choice of these connectivity densities mirrors previous work [22].

Synapses are modeled in the double exponential formalism:

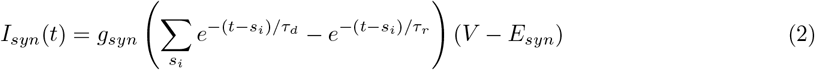

*g*_*syn*_ is the maximum conductance of the synapse, *V* the membrane voltage of the post-synaptic neuron, *E*_*syn*_ the synaptic current’s reversal potential, *τ*_*d*_ and *τ*_*r*_ the synaptic decay and rise time constants, and *s*_*i*_ represents the times of all pre-synaptic spikes occurring before the current time *t. g*_*syn*_ = 0.00235 mS/cm^2^ for E-I synapses, 0.003 mS/cm^2^ for I-E synapses, and 0.025 mS/cm^2^ for I-I synapses. For excitatory synapses, *E*_*syn*_ = 0 mV, *τ*_*d*_ = 3.0 ms, and *τ*_*r*_ = 0.2 ms. For inhibitory synapses, *E*_*syn*_ = −75 mV as a default (but is varied throughout this work), *τ*_*d*_ = 5.5 ms, and *τ*_*r*_ = 0.2 ms. These parameter values have been previously shown to yield classical “weak PING” dynamics [22].

### Noisy Input

External noisy input to the network is simulated via an independent Ornstein–Uhlenbeck process [73–75] to each neuron. At time *t* the noisy input *I*_*noise*_(*t*) is defined as

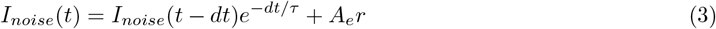

where *dt* is the integration step, *τ* = 5 ms, and *r* is a random number from the standard normal distribution. The constant 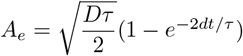, where *D* determines the noise amplitude.

Rather than reporting the noise amplitudes *D*, we instead report the consequent fluctuations in membrane potential. This was determined by holding a neuron below the spike threshold with a tonic current input of −0.4 *µ*A/cm^2^, stimulating with noisy input for 50000 ms, and calculating the standard deviation of the voltage (*SD*_*V*_) over this period.

### Initial Conditions

Each neuron is initialized with variability in *V, h*, and *n*. Under the default, “highly active” initial conditions, each neuron receives its defined *I*_*ext*_ from *t* = 0 ms, but spikes do not trigger synaptic signaling until after *t* = 250 ms to allow initial transients to decay. For the altered “sparsely active” initial conditions, *I*_*ext*_ for E cells is altered for *t <* 2000 so that these values are randomly sampled from a uniform distribution [−0.06, −0.04] *µ*A/cm^2^ before reverting to the default values defined previously at *t* = 2000 ms. Additionally, synaptic signalling is only triggered by spikes occurring after *t* = 1000 ms.

### Numerics

Simulations were performed in Python 3 using NumPy, utilizing Linux-based high performance computing resources provided by SickKids and UConn. Equations are integrated using the Euler-Maruyama method, with a time step *dt* = 0.02 ms and a simulation duration of 20,000 ms throughout.

### Measure and Quantifications

Firing rate histograms were generated by simply convolving each spike with a Gaussian kernel with a standard deviation of 2 ms. “Inconsistent oscillatory activity” was identified if a measure of network synchrony (used throughout the authors’ previous work [22, 23, 69, 76]), calculated in moving time windows fell below a threshold value of 0.2. In Figure 4 the existence of stable oscillations was calculated by detecting whether the peaks of the excitatory firing rate histogram ever exceeded 40. Thorough inspection confirmed these parallel mechanisms of detecting synchronous network oscillations were equivalent.

Network oscillation frequency was calculated via the peaks in a firing rate histogram equivalent used as part of the calculation of network synchrony cited above. These peaks were thresholded so that the frequency was only influenced by periods in which the network was clearly oscillatory. Neuron spike rate represents the mean spiking rate of each neuron in a given population, calculated simply by summing the number of spikes and dividing by the number of cells and duration of interest.

When transient periods of oscillatory activity with varying strengths were qualitatively observed in systems not exhibiting PING-like oscillations, we quantitatively deemed such activity a “gamma event” through distinguishing features of the inhibitory firing rate histogram. A gamma event was detected if ≥ 3 subsequent peaks of the inhibitory firing rate histogram occurred at at least 7 mean inhibitory spikes per cell per second. The examples in Figure 4**B-C** are centered on these events, and a horizontal line demarcating this threshold is included for reference.

To confirm that these events corresponded with gamma activity, we calculated spectograms of the excitatory firing rate histograms using the SciPy Python package. This was done on firing rate histograms with additional temporal precision so that a sample is taken at intervals equal to *dt*. The *nperseg* parameter is set equal to 6400 to achieve temporal discretization in segments of approximately 100 ms and spectral discretization in segments of approximately 10 Hz. To specifically quantify gamma activity, the power at frequencies between 30 and 80 are summed to create a time series of “gamma power” (the bottom-right panels in Figure 4**B-C**). Thorough inspection confirmed that the events distinguished by the features of the I cell firing rate histograms described above also yielded notable increases in gamma power in the E cell firing rate histogram, justifying classifying them as gamma events.

The duration of gamma events was the difference between the timing of the first and last peak in the I cell firing rate histogram demarcating the event. The peak frequency of the gamma events was the frequency exhibiting maximal power in the spectogram analysis of the E cell firing rate histogram during this period.

### Mathematical Analyses

To investigate whether oscillatory dynamics analogous to those observed in our spiking neuronal network might arise from a subcritical Hopf bifurcation in an idealized system, we studied a model exhibiting such structure as defined by the following polar equations [49]:

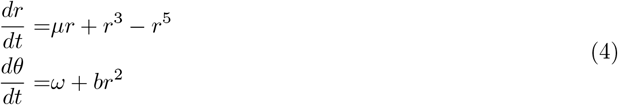

Here, *r* is the radius and *θ* is the phase. *ω* = 0.5 in all settings. Figure 5**A** has *µ* = −0.245, Figure 5**B** has *µ* = 0.1, and Figure 5**C** has *µ* = −0.255. To facilitate visualization the phase portraits were generated with *b* = 2 while the activity traces were generated with *b* = 6 (*b* does not affect the system’s stability structure). Euler integration was used to numerically solve the system with time step *dt* = 0.001 ms. The position of the unstable limit cycle was directly determined by solving for an unstable solution to *dr/dt* = 0. The two stable solutions (fixed point and limit cycle) are apparent both analytically and numerically when creating the phase portrait. In the bottom activity traces, synthetic perturbations were introduced at 0, 50, and 100 ms to assess oscillatory features arising from varying perturbation magnitudes. Each perturbation lasted for 1 ms, setting *r* = 0.3, 0.6, and 0.9 respectively.

We also utilized the Wilson-Cowan framework to assess transient oscillations in a more biologically motivated system. The exact version of the equations implemented are as follows [77]:

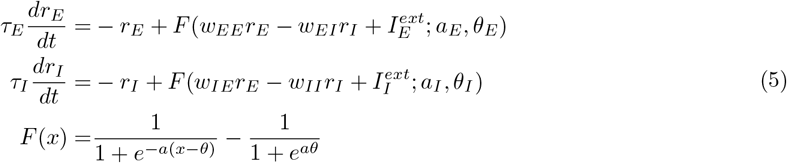

Parameters were derived from bifurcation analyses performed in previous literature [77]: *τ*_*E*_ = 20 ms, *τ*_*I*_ = 10 ms, *a*_*E*_ = *a*_*I*_ = 1.0, *θ*_*E*_ = 5.0, *θ*_*I*_ = 20.0, *w*_*EE*_ = 16.0, *w*_*EI*_ = 26.0, *w*_*E*_ = 20.0, *w*_*II*_ = 0.5, 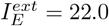, and 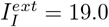.

To approximate realistic fluctuations in neural population activity (Figure 5**D**, right), Ornstein-Uhlenbeck noise was added to both terms and integrated using the Euler-Murayama technique as previously described for the spiking neuronal network. The parameters used were *D* = 5 ∗ 10^−7^, *τ* = 5 ms, and *dt* = 0.1 ms. Spectograms were computed from the resulting time-series data using SciPy functions as previously described.

### Significance testing

All significance tests (see Figure 4) were performed using functions from the SciPy Python package. A standard threshold of p *<* 0.05 is used to report statistically significant differences.

## Code Accessibility

The model code will be made openly accessible via a GitHub repository upon final publication of this manuscript.

## Acknowledgements

Funding for this research provided by CIHR Foundation Grant 167276 (S.A.P.) and a CIHR Postdoctoral Fellowship (S.R.).

## References

[1] Buzsaki, G. Rhythms of the Brain (Oxford university press, 2006).

[2] Buzsaki, G. & Draguhn, A. Neuronal oscillations in cortical networks. Science 304, 1926–1929 (2004).

[3] Buzsáki, G. & Wang, X.-J. Mechanisms of gamma oscillations. Annual Review of Neuroscience 35, 203–225 (2012).

[4] Ahmed, O. J. & Cash, S. S. Finding synchrony in the desynchronized eeg: the history and interpretation of gamma rhythms. Frontiers in Integrative Neuroscience 7, 58 (2013).

[5] Han, C., Shapley, R. & Xing, D. Gamma rhythms in the visual cortex: functions and mechanisms. Cognitive Neurodynamics 16, 745–756 (2022).

[6] Jia, X. & Kohn, A. Gamma rhythms in the brain. PLoS Biology 9, e1001045 (2011).

[7] Buzsáki, G. & Moser, E. I. Memory, navigation and theta rhythm in the hippocampal-entorhinal system. Nature neuroscience 16, 130–138 (2013).

[8] Kucewicz, M. T. et al. Dissecting gamma frequency activity during human memory processing. Brain 140, 1337–1350 (2017).

[9] Womelsdorf, T. & Fries, P. The role of neuronal synchronization in selective attention. Current Opinion in Neurobiology 17, 154–160 (2007).

[10] Ashwin, P., Coombes, S. & Nicks, R. Mathematical frameworks for oscillatory network dynamics in neuroscience. The Journal of Mathematical Neuroscience 6, 1–92 (2016).

[11] Gray, C. M. Synchronous oscillations in neuronal systems: mechanisms and functions. Journal of Computational Neuroscience 1, 11–38 (1994).

[12] Traub, R. D., Jefferys, J. G. & Whittington, M. A. Simulation of gamma rhythms in networks of interneurons and pyramidal cells. Journal of Computational Neuroscience 4, 141–150 (1997).

[13] Whittington, M. A., Traub, R. D. & Jefferys, J. G. Synchronized oscillations in interneuron networks driven by metabotropic glutamate receptor activation. Nature 373, 612–615 (1995).

[14] Ermentrout, G. B. & Kopell, N. Fine structure of neural spiking and synchronization in the presence of conduction delays. Proceedings of the National Academy of Sciences 95, 1259–1264 (1998).

[15] Whittington, M. A., Traub, R. D., Kopell, N., Ermentrout, B. & Buhl, E. H. Inhibition-based rhythms: experimental and mathematical observations on network dynamics. International Journal of Psychophysiology 38, 315–336 (2000).

[16] Cowan, J. D., Neuman, J. & van Drongelen, W. Wilson–cowan equations for neocortical dynamics. The Journal of Mathematical Neuroscience 6, 1 (2016).

[17] Börgers, C. & Kopell, N. Effects of noisy drive on rhythms in networks of excitatory and inhibitory neurons. Neural Computation 17, 557–608 (2005).

[18] Keeley, S., Byrne, Á., Fenton, A. & Rinzel, J. Firing rate models for gamma oscillations. Journal of Neurophysiology 121, 2181–2190 (2019).

[19] Börgers, C., Talei Franzesi, G., LeBeau, F. E., Boyden, E. S. & Kopell, N. J. Minimal size of cell assemblies coordinated by gamma oscillations. PLoS Computational Biology 8, e1002362 (2012).

[20] Börgers, C. & Kopell, N. Synchronization in networks of excitatory and inhibitory neurons with sparse, random connectivity. Neural Computation 15, 509–538 (2003).

[21] Börgers, C. & Walker, B. Toggling between gamma-frequency activity and suppression of cell assemblies. Frontiers in Computational Neuroscience 7, 33 (2013).

[22] Rich, S., Zochowski, M. & Booth, V. Dichotomous dynamics in ei networks with strongly and weakly intra-connected inhibitory neurons. Frontiers in Neural Circuits 11, 104 (2017).

[23] Rich, S., Zochowski, M. & Booth, V. Effects of neuromodulation on excitatory–inhibitory neural network dynamics depend on network connectivity structure. Journal of Nonlinear Science 30, 1–24 (2018).

[24] Olufsen, M. S., Whittington, M. A., Camperi, M. & Kopell, N. New roles for the gamma rhythm: population tuning and preprocessing for the beta rhythm. Journal of Computational Neuroscience 14, 33–54 (2003).

[25] Krupa, M., Gielen, S. & Gutkin, B. Adaptation and shunting inhibition leads to pyramidal/interneuron gamma with sparse firing of pyramidal cells. Journal of Computational Neuroscience 37, 357–376 (2014).

[26] Almog, M. & Korngreen, A. Is realistic neuronal modeling realistic? Journal of Neurophysiology 116, 2180–2209 (2016).

[27] Fernandez-Ruiz, A., Sirota, A., Lopes-dos Santos, V. & Dupret, D. Over and above frequency: gamma oscillations as units of neural circuit operations. Neuron 111, 936–953 (2023).

[28] Brunel, N. & Wang, X.-J. What determines the frequency of fast network oscillations with irregular neural discharges? i. synaptic dynamics and excitation-inhibition balance. Journal of Neurophysiology 90, 415–430 (2003).

[29] Dubey, A. & Ray, S. Comparison of tuning properties of gamma and high-gamma power in local field potential (lfp) versus electrocorticogram (ecog) in visual cortex. Scientific Reports 10, 5422 (2020).

[30] Tahvili, F., Vinck, M. & di Volo, M. Pv and som cells play distinct causal roles in controlling network oscillations and stability. Cell Reports 44 (2025).

[31] McCormick, D. A. Gaba as an inhibitory neurotransmitter in human cerebral cortex. Journal of Neurophysiology 62, 1018–1027 (1989).

[32] Deco, G., Jirsa, V., McIntosh, A. R., Sporns, O. & Kötter, R. Key role of coupling, delay, and noise in resting brain fluctuations. Proceedings of the National Academy of Sciences 106, 10302–10307 (2009).

[33] Mori, T. & Kai, S. Noise-induced entrainment and stochastic resonance in human brain waves. Physical Review Letters 88, 218101 (2002).

[34] Fraiman, D. & Chialvo, D. R. What kind of noise is brain noise: anomalous scaling behavior of the resting brain activity fluctuations. Frontiers in Physiology 3, 307 (2012).

[35] Bernasconi, F. et al. Noise in brain activity engenders perception and influences discrimination sensitivity. Journal of Neuroscience 31, 17971–17981 (2011).

[36] Jeong, H. Y. & Gutkin, B. Synchrony of neuronal oscillations controlled by gabaergic reversal potentials. Neural Computation 19, 706–729 (2007).

[37] Barmashenko, G., Hefft, S., Aertsen, A., Kirschstein, T. & Köhling, R. Positive shifts of the gabaa receptor reversal potential due to altered chloride homeostasis is widespread after status epilepticus. Epilepsia 52, 1570–1578 (2011).

[38] Palma, E. et al. Anomalous levels of cl-transporters in the hippocampal subiculum from temporal lobe epilepsy patients make gaba excitatory. Proceedings of the National Academy of Sciences 103, 8465–8468 (2006).

[39] Miles, R., Blaesse, P., Huberfeld, G., Wittner, L. & Kaila, K. Chloride homeostasis and gaba signaling in temporal lobe epilepsy. Jasper’s Basic Mechanisms of the Epilepsies, Fourth Edition (2012).

[40] Price, T. J., Cervero, F., Gold, M. S., Hammond, D. L. & Prescott, S. A. Chloride regulation in the pain pathway. Brain Research Reviews 60, 149–170 (2009).

[41] Coull, J. A. et al. Trans-synaptic shift in anion gradient in spinal lamina i neurons as a mechanism of neuropathic pain. Nature 424, 938–942 (2003).

[42] Cherubini, E., Di Cristo, G. & Avoli, M. Dysregulation of gabaergic signaling in neurodevelomental disorders: targeting cation-chloride co-transporters to re-establish a proper e/i balance. Frontiers in Cellular Neuroscience 15, 813441 (2022).

[43] Ben-Ari, Y. Excitatory actions of gaba during development: the nature of the nurture. Nature Reviews Neuroscience 3, 728–739 (2002).

[44] Currin, C. B. & Raimondo, J. V. Computational models reveal how chloride dynamics determine the optimal distribution of inhibitory synapses to minimise dendritic excitability. PLoS Computational Biology 18, e1010534 (2022).

[45] Doyon, N. et al. Efficacy of synaptic inhibition depends on multiple, dynamically interacting mechanisms implicated in chloride homeostasis. PLoS Computational Biology 7, e1002149 (2011).

[46] Doyon, N., Vinay, L., Prescott, S. A. & De Koninck, Y. Chloride regulation: a dynamic equilibrium crucial for synaptic inhibition. Neuron 89, 1157–1172 (2016).

[47] Jedlicka, P. & Backus, K. Inhibitory transmission, activity-dependent ionic changes and neuronal network oscillations. Physiological Research 55, 139 (2006).

[48] Wilson, H. R. & Cowan, J. D. Excitatory and inhibitory interactions in localized populations of model neurons. Biophysical journal 12, 1–24 (1972).

[49] Strogatz, S. H. Nonlinear Dynamics and Chaos: With Applications to Physics, Biology, Chemistry, and Engineering (Chapman and Hall/CRC, 2024).

[50] Jiruska, P. et al. Synchronization and desynchronization in epilepsy: controversies and hypotheses. The Journal of Physiology 591, 787–797 (2013).

[51] Börgers, C. Weak ping rhythms. An Introduction to Modeling Neuronal Dynamics 281–292 (2017).

[52] Hiver, P. Attractor states. Motivational Dynamics in Language Learning 20–28 (2015).

[53] Kopell, N., Börgers, C., Pervouchine, D., Malerba, P. & Tort, A. Gamma and theta rhythms in biophysical models of hippocampal circuits. Hippocampal Microcircuits 423–457 (2010).

[54] Aghamohammadi, C., Chandrasekaran, C. & Engel, T. A. A doubly stochastic renewal framework for partitioning spiking variability. Nature communications 16, 8656 (2025).

[55] Veselić, K. Damped oscillations of linear systems: a mathematical introduction, vol. 2023 (Springer Science & Business Media, 2011).

[56] Destexhe, A. & Rudolph-Lilith, M. Neuronal noise, vol. 8 (Springer Science & Business Media, 2012).

[57] Matthews, P. Relationship of firing intervals of human motor units to the trajectory of post-spike after-hyperpolarization and synaptic noise. The Journal of Physiology 492, 597–628 (1996).

[58] Wagner, S., Castel, M., Gainer, H. & Yarom, Y. Gaba in the mammalian suprachiasmatic nucleus and its role in diurnal rhythmicity. Nature 387, 598–603 (1997).

[59] Tian, J.-b. et al. Trpc4 and girk channels underlie neuronal coding of firing patterns that reflect gq/11–gi/o coincidence signals of variable strengths. Proceedings of the National Academy of Sciences 119, e2120870119 (2022).

[60] Renart, A. et al. The asynchronous state in cortical circuits. Science 327, 587–590 (2010).

[61] Börgers, C., Krupa, M. & Gielen, S. The response of a classical hodgkin–huxley neuron to an inhibitory input pulse. Journal of Computational Neuroscience 28, 509–526 (2010).

[62] Meng, J. H. & Riecke, H. Synchronization by uncorrelated noise: interacting rhythms in interconnected oscillator networks. Scientific Reports 8, 6949 (2018).

[63] Cembrowski, M. S. & Menon, V. Continuous variation within cell types of the nervous system. Trends in Neurosciences 41, 337–348 (2018).

[64] Cembrowski, M. S. & Spruston, N. Heterogeneity within classical cell types is the rule: lessons from hippocampal pyramidal neurons. Nature Reviews Neuroscience 20, 193–204 (2019).

[65] Scala, F. et al. Phenotypic variation of transcriptomic cell types in mouse motor cortex. Nature 598, 144–150 (2021).

[66] Zeng, H. & Sanes, J. R. Neuronal cell-type classification: challenges, opportunities and the path forward. Nature Reviews Neuroscience 18, 530–546 (2017).

[67] Zeng, H. What is a cell type and how to define it? Cell 185, 2739–2755 (2022).

[68] Tasic, B. et al. Shared and distinct transcriptomic cell types across neocortical areas. Nature 563, 72–78 (2018).

[69] Rich, S., Chameh, H. M., Lefebvre, J. & Valiante, T. A. Loss of neuronal heterogeneity in epileptogenic human tissue impairs network resilience to sudden changes in synchrony. Cell Reports 39 (2022).

[70] Hutt, A., Rich, S., Valiante, T. A. & Lefebvre, J. Intrinsic neural diversity quenches the dynamic volatility of neural networks. Proceedings of the National Academy of Sciences 120, e2218841120 (2023).

[71] Fink, C. G., Booth, V. & Zochowski, M. Cellularly-driven differences in network synchronization propensity are differentially modulated by firing frequency. PLoS Computational Biology 7, e1002062 (2011).

[72] Stiefel, K. M., Gutkin, B. S. & Sejnowski, T. J. Cholinergic neuromodulation changes phase response curve shape and type in cortical pyramidal neurons. PloS One 3, e3947 (2008).

[73] Uhlenbeck, G. E. & Ornstein, L. S. On the theory of the brownian motion. Physical Review 36, 823 (1930).

[74] Destexhe, A., Rudolph, M.Fellous, J.-M. & Sejnowski, T. J. Fluctuating synaptic conductances recreate in vivo-like activity in neocortical neurons. Neuroscience 107, 13–24 (2001).

[75] Piwkowska, Z. et al. Characterizing synaptic conductance fluctuations in cortical neurons and their influence on spike generation. Journal of Neuroscience Methods 169, 302–322 (2008).

[76] Rich, S. et al. Inhibitory network bistability explains increased interneuronal activity prior to seizure onset. Frontiers in Neural Circuits 13, 81 (2020).

[77] Li, X., Li, Z., Yang, W., Wu, Z. & Wang, J. Bidirectionally regulating gamma oscillations in wilson-cowan model by self-feedback loops: A computational study. Frontiers in Systems Neuroscience 16, 723237 (2022).

