## Supplemental Materials for "Physiologically realistic gamma activity produced *in silico* by weakening the PING attractor state"

**This PDF file includes:**

Fig. S1
Legends for Movies S1 to S4

**Other supporting materials for this manuscript include the following:**

Movies S1 to S4

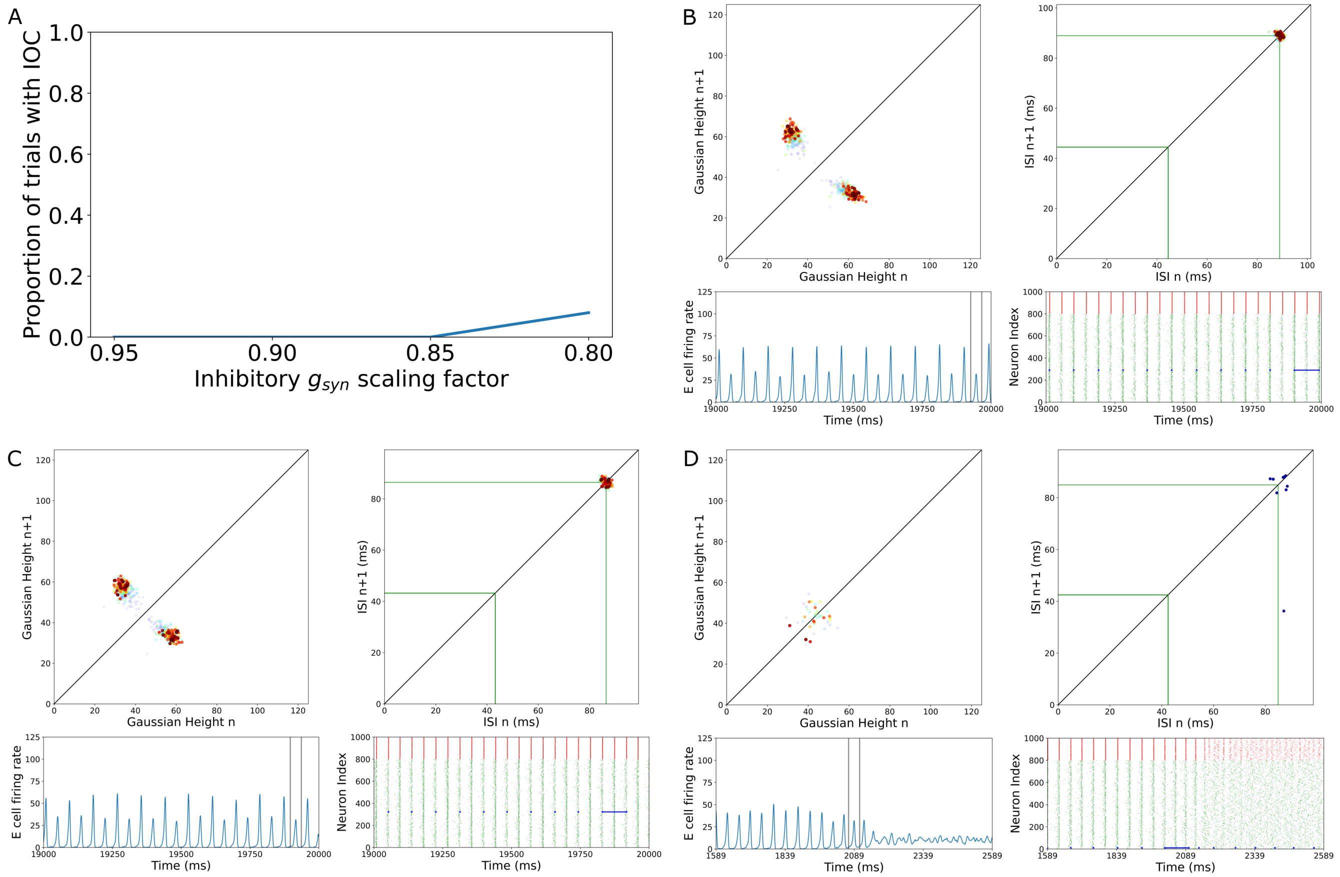


**Fig. S1. Downscaling inhibitory synaptic weights does not reproduce dynamics of depolarized E$_{GABA}$. A**: Proportion of 100 independent simulations for E$_{GABA}$=-75 mV that exhibit inconsistent oscillatory activity (IOC) with decreased inhibitory synaptic weights ($g_{syn} = (\text{Scaling Factor}) * g_{syn}^{default}$ for both inhibitory synapses with $g_{syn}^{default}$ as reported in the Materials and Methods). **B-D**: Example dynamics with a scaling factor of 0.85 (Panel **B**) and 0.8 (Panels **C-D**), with panels as in Figure 3. Dynamics in Panels **B** and **C** are qualitatively similar to those in the default network with E$_{GABA}$=-75 mV (Figure 1**A**). While a small proportion of simulations with a scaling factor of 0.80 will exhibit a breakdown of oscillatory activity, as exemplified in Panel **D**, in all cases this breakdown occurs very early in the simulation. Furthermore, transitions into and out of synchrony (Figure 2**A**) that distinguish systems with sufficiently depolarized E$_{GABA}$ never arise.

**Movie S1.** ***Dynamics of system in Figure 3D visualized over time.*** **Panels as in Figure 3: Pointcare return maps (top left) of continuous firing rate histogram peaks (bottom left), and Pointcare return maps (top right) of an individual cell's inter-spike intervals (highlighted in blue on the raster plot in the bottom right), with gray lines representing integer multiples of the network oscillatory period. As the return maps are generated over time, new data points are plotted with no opacity while older points fade, and colors shift from blue to red as the simulation progresses over 20,000 ms. Vertical lines in bottom-left surround the peak in the firing rate histogram most recently plotted; horizontal blue line in bottom-right connects the spikes of the chosen cell generating the inter-spike interval most recently plotted.**

**Movie S2.** ***Dynamics of system in Figure 3E visualized over time.*** **Panels and plotting as described in Movie S1.**

**Movie S3.** ***Dynamics of system in Figure 3F visualized over time.*** **Panels and plotting as described in Movie S1.**

**Movie S4.** ***Dynamics of system in Figure 3G visualized over time.*** **Panels and plotting as described in Movie S1. Movie stops when oscillatory activity terminates in the system.**
